# MX2 Mediates Collapse of the HIV-1 Capsid

**DOI:** 10.64898/2026.05.07.723526

**Authors:** Andrew Goodale, Szu-Wei Huang, Nathalia Almeida, Dave J. Williamson, Nikoloz Shkriabai, Gilberto Betancor, Luis Apolonia, Frank DiMaio, Sergi Padilla-Parra, Mamuka Kvaratskhelia, Julien R. C. Bergeron, Michael H. Malim

## Abstract

The HIV-1 capsid core encapsulates the viral genome and mediates its delivery into the host cell’s nucleus. It is composed of multiple copies of the Capsid (CA, p24^Gag^) protein, assembled into hexamers and pentamers to create a lattice that forms a fullerene-like cone. Myxovirus resistance 2 (MX2) is an HIV-1 restriction factor that binds to the capsid core and blocks nuclear import of the viral genome. Here, we define a minimal region of MX2 required for HIV-1 restriction and produce a corresponding functional recombinant protein. We have used cryo-electron microscopy to determine the structure of this MX2 fragment bound to the tri-hexamer interface of the capsid lattice, revealing a large, buried interface combining electrostatic and hydrophobic interactions. This structure, together with assays that measure capsid core destabilisation, shows that MX2 binding induces conformational rearrangements in the capsid lattice that culminate in a loss of integrity. These results support a model whereby MX2 exerts its antiviral activity by disrupting the capsid lattice, inducing premature fragmentation and preventing HIV-1 nuclear import. By revealing the structural basis for MX2-mediated restriction, this work also provides the framework for the development of anti-HIV molecules that mimic MX2 restriction.

## Main

The HIV-1 capsid core encapsulates the viral genome and proteins essential for replication, and protects them from innate immune sensors^1,2^. It is composed of ∼1500 copies of the viral CA protein, assembled into ∼250 hexamers and 12 pentamers, to form a fullerene-like cone^3^ that transports the HIV-1 genome from the plasma membrane to the nucleus. This process relies on interactions with various host cell cofactors, which contribute to capsid stability^4^, nuclear import^5–7^ and integration^8^.

Myxovirus resistance 2 (MX2, also referred to as MXB) is an interferon inducible restriction factor of HIV-1^9,10^ as well as herpesviruses^11^, hepatitis B virus^12^ and hepatitis C virus^13^. MX2 inhibits HIV-1 infection post-entry by binding to the HIV-1 capsid and blocking viral nuclear import. MX2 belongs to a family of dynamin-like GTPases^14^ and is comprised of a disordered N-terminal domain (NTD) followed by a GTPase domain and an extended stalk domain that are separated by a tripartite bundle signalling element (BSE). Like other dynamin-like GTPases^15^, MX2 forms antiparallel dimers^16^ that can assemble into higher-order oligomers in a GTP hydrolysis-dependent manner^17^. GTPase activity is required for HSV inhibition^18^ but is dispensable for HIV-1 restriction, which is instead mediated by the NTD^19^, and transfer of the MX2 NTD to unrelated scadold proteins is sudicient to confer HIV-1 inhibition^19^.

The NTD is responsible for the localisation of MX2 to the cytoplasmic side of the nuclear envelope and contains a conserved triple arginine motif (residues 11-13, Fig. 1a) that is essential for its antiviral activity^19^, capsid binding^20^ and interactions with host factors such as nucleoporins^21^ and components of the myosin light chain phosphatase (MLCP)^22^. MLCP regulates MX2 activity through serine phosphorylation at multiple sites within the NTD which can either enhance or supress MX2’s antiviral function^22,23^. Molecular dynamic (MD) simulations have predicted that the NTD of MX2 engages the capsid lattice through electrostatic interactions between the MX2 11-13 RRR motif and the negatively charged capsid tri-hexamer interface^24^, although the molecular basis for their interaction and the consequences for lattice structure and geometry remain poorly understood. Notably, several CA mutations have been reported that reduce or escape MX2 inhibition, including mutations within the cyclophilin A (CypA) binding loop, the phenylalanine-glycine (FG) dipeptide repeat binding pocket, and at the tri-hexamer interface^9,10,24,25^.

**Figure 1:**
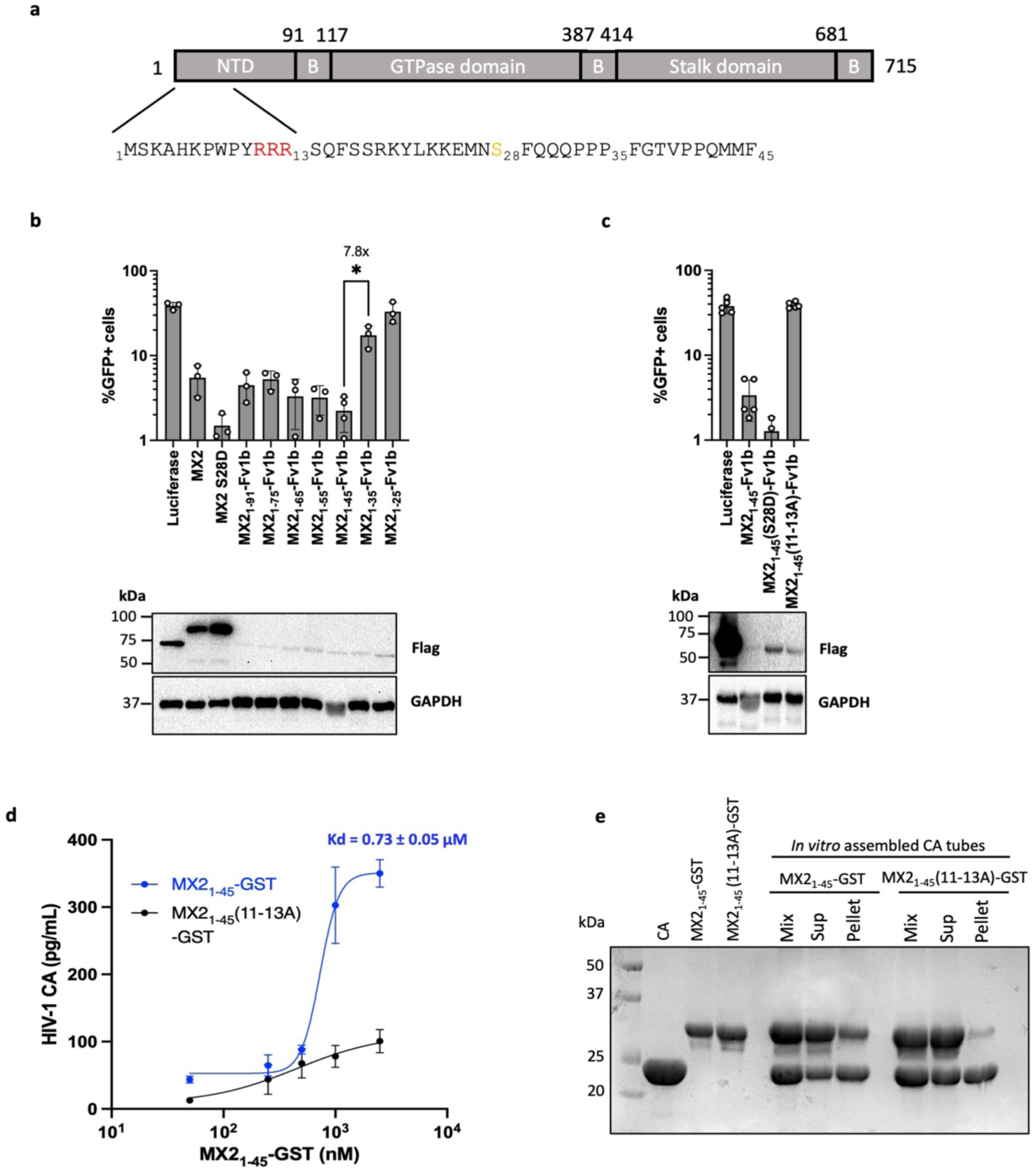
Minimal MX2 fragment retains HIV-1 restriction and binds to capsid. **a,** Domain organisation of MX2 showing residues 1–45, with the 11–13 RRR motif (red) and S28 (yellow) indicated. **b,** U87-MG cells were transduced with EasiLV particles encoding FLAG-tagged luciferase, wild-type or mutant MX2, or MX2_NTD_–Fv1b fusion proteins. HIV-1/GFP infectivity was assessed 48 h post-infection by flow cytometry. Data represent mean ± SD from three independent biological replicates. Protein expression was confirmed by immunoblot (below). Two-tailed paired t-test, * p<0.05. **c,** U87-MG cells expressing luciferase or MX2_1–45_–Fv1b variants were challenged with HIV-1/GFP, and the proportion of GFP⁺ cells was quantified by flow cytometry. Data represent mean ± SD from three independent biological replicates. Two-tailed paired t-test, * p<0.05. Protein expression was confirmed by immunoblot (below). **d,** Quantitation of GST-mediated affinity pull-down of native HIV-1 cores bound to indicated concentrations of MX2_1-45_-GST or MX2_1–45_(RRR/AAA)-GST. Each data point represents mean values ± SD from two independent biological replicates. **e,** Pelleting assay of MX2_1-45_-GST or MX2_1–45_(RRR/AAA)-GST incubated with capsid tubes (Total) prior to centrifugation to separate the soluble (Sup) and insoluble (Pellet) fractions.

Here, we identify a minimal functional region of the MX2 NTD, and report the purification of a functional protein encompassing this region. Using cryogenic electron microscopy (cryo-EM), we have determined the structure of the HIV-1 capsid lattice in complex with this MX2 NTD fragment. This reveals the molecular details of the MX2 binding pocket at the CA tri-hexamer interface, and demonstrates that MX2 binding induces marked conformational changes within the lattice that result in capsid lattice remodelling and unprogrammed fragmentation.

### A minimal MX2 construct restricts HIV-1 infection and binds to the capsid core

To help define the functional boundaries of the MX2 NTD, we engineered a set of truncated variants and determined their capacity to restrict HIV-1 infection. Specifically, MX2_NTD_-Fv1b chimeras with various MX2 NTD boundaries were expressed within U87-MG cells and infected with a HIV-1 based lentiviral vector encoding GFP as a reporter (HIV-1-GFP) (Extended Data Fig. 1). Consistent with previous data^10,19,22,23^, cells expressing wild type MX2 showed ∼10-fold lower levels of GFP expression compared to the luciferase control (Fig. 1b). MX2_1-45_-Fv1b chimeras containing MX2 residues 1-45 (as well as longer constructs) retained full antiviral activity when compared to full length MX2. In contrast, both MX2_1-35_-Fv1b and MX2_1-25_-Fv1b lost their ability to restrict HIV-1 (Fig. 1b). As expected, the MX2_1-45_-Fv1b R11A/R12A/R13A mutant (hereafter referred to as MX2_1–45_(RRR/AAA)) abolished antiviral activity^19^. Furthermore, the MX2_1-45_-Fv1b construct with the S28D mutation (Fig. 1a), which mimics constitutive phosphorylation and enhances MX2 antiviral activity^23^, also displayed increased activity further validating the utility of studying the N-terminal 1-45 fragment (Fig. 1c).

To explore the physical interaction between MX2 and CA, we engineered and purified a chimeric protein comprising the N-terminal 45 amino acids of MX2 fused to the N-terminus of the dimeric protein GST (MX2_1-45_-GST). We first tested whether MX2_1-45_-GST binds to CA in the context of native capsid cores isolated from HIV-1 virions (Fig. 1d). MX2-bound CA was captured via glutathione binding and then quantified by ELISA, revealing a binding affinity of 0.74 µM. Binding to the capsid lattice was then confirmed by mixing MX2_1-45_-GST with *in vitro* assembled CA tubes followed by co-pelleting using centrifugation (Fig. 1e). Importantly, the specificity of both binding reactions was demonstrated by the lack of binding of MX2_1-45_-GST carrying the inactivating RRR mutation.

### MX2 induces capsid lattice collapse

Inositol hexakisphosphate (IP6) is a host cofactor that facilitates HIV-1 maturation and stabilises the capsid lattice by binding to a ring Arg-18 and Lys-25 residues at the central pore of capsid hexamers and pentamers^26–28^. IP6 also induces assembly of the capsid lattice *in vitro*^29^, with IP6-mediated assembly at pH 8.0 resulting in capsid morphologies ranging from capped tubes to cones (Extended Data Fig. 2a); whereas at pH 6.0, IP6 facilitates the formation of conically shaped capsid like particles (CLPs) (Fig. 2a), which more closely mimic native viral cores.

**Figure 2:**
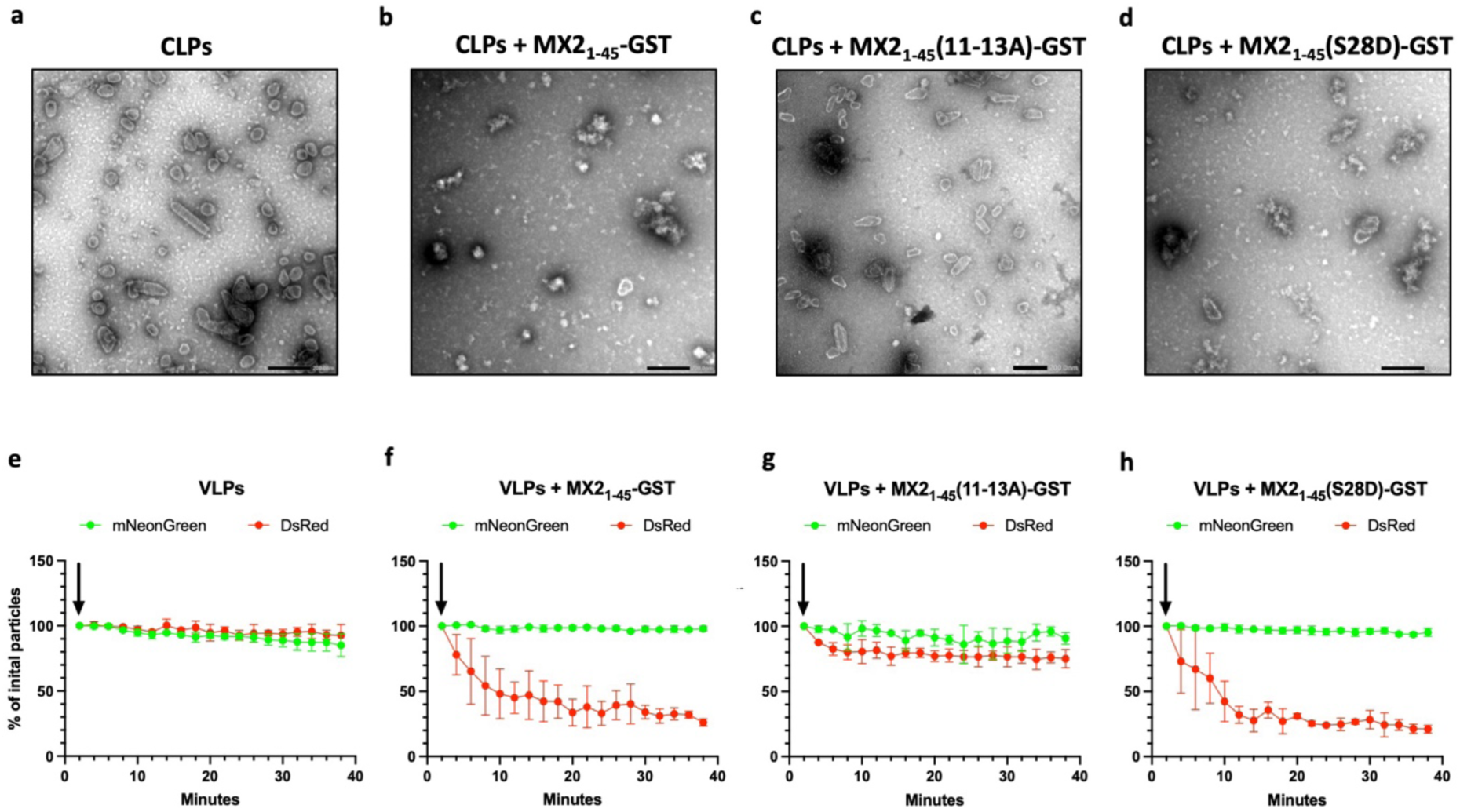
MX2 disrupts the integrity of the capsid lattice. **a-d**, Negative stain TEM micrographs of CLPs assembled from CA in 500 µM IP6, 50 mM MES pH 6.0, and 150 mM NaCl in the presence or absence of the indicated MX2_1–45_-GST constructs. Scale bars, 200 nm. **e-f,** IN-mNeonGreen and CypA-DsRed-labelled virus like particles (VLPs) were adhered to coverslips and permeabilisation buffer containing 50 µg mL−1 saponin, 20 µM IP6 and the indicated the MX2_1-45_-GST constructs. Fluorescence intensities of mNeonGreen and DsRed were quantified over time by threshold-based particle detection, and the number of fluorescent particles above threshold was normalised to the initial time point. Arrows indicate the time at which the permeabilisation buffer was added.

To assess whether and how MX2 binding impacts the morphology of capsid tubes or CLPs, capsid complexes assembled with IP6 at pH 6.0 were incubated with MX21–45–GST, MX2_1–45_(RRR/AAA)-GST or MX2_1–45_(S28D)-GST and imaged by negative-stain transmission electron microscopy (TEM). MX2-bound capsid tubes appeared heavily decorated, losing their defined edges, and bundled into irregular clusters. Nevertheless, the tubes remained mostly intact, and contained capped ends, indicating that pentamers remained incorporated into the MX2–capsid complexes (Extended Data Fig. 2). In sharp contrast, when MX2_1–45_–GST or MX2_1–45_(S28D)-GST were incubated with CLPs, a profound collapse of CLP conical morphology was evident together with the extensive and disordered aggregation of material (Fig. 2b, Fig. 2d). By contrast, CLPs remained intact when incubated with the MX2_1–45_(RRR/AAA)-GST (Fig. 2c), indicating that CLP destruction is triggered by MX2 binding. These observations indicate that MX2 binding perturbs capsid lattice integrity, and show that the destabilising consequences are morphologically more severe for CLPs than for tubes.

To address whether the destabilisation reconstituted CLPs mirrors MX2’s effects on viral particles produced from human cells, we measured lattice integrity using a fluorescence-based assay where virions were co-labelled with integrase (IN)-mNeonGreen and CypA-DsRed^30^. Virus particles were immobilised on coverslips and imaged to quantify the co-localisation of the two fluorophores. A reduction in CypA-DsRed signal over time relative to IN-mNeonGreen indicates capsid disassembly, allowing us to quantify the impact of MX2 binding on core stability for individual virions.

Most CypA-DsRed puncta remain co-localised with IN-mNeonGreen following viral membrane permeabilisation (Fig. 2e), consistent with previous reports^26^. We found that CypA-DsRed puncta were substantially reduced when viral cores were incubated with MX2_1–45_–GST and that the rate of loss was further accelerated using the phosphomimetic gain-of-function mutant, MX2_1–45_(S28D)-GST (Fig. 2f,h, Extended Data Fig. 3). As expected, the inactive MX2_1–45_(RRR/AAA)-GST mutant had minimal edect (Fig. 2g), confirming the importance of the MX2 RRR motif for its interaction with the capsid lattice. We note that it is possible that the loss of red puncta could indicate MX2 outcompeting CypA for capsid binding; however, the CypA binding site does not overlap with that of MX2^4,31^. Together, these observations reveal that MX2 binding to the capsid lattice results in the destruction of HIV-1 cores.

### Structural basis for the MX2-mediated capsid destabilisation

We next used cryo-electron microscopy (cryo-EM) to characterise the structure of MX2 in complex with the capsid lattice. Given that the incubation of CLPs with MX2 resulted in the dramatic loss of capsid structure, CA was pre-assembled into tubes prior to MX2 addition (See above, Extended Data Fig. 2). We also utilised the gain-of-function mutant MX2_1–45_(S28D)-GST for these structural studies, reasoning that the S28D mutation may increase binding adinity to the lattice and consequently reduce structural heterogeneity, making it more amenable to high resolution analysis.

Cryo-EM datasets were collected for IP6-stabilised capsid tubes alone and in complex with MX2_1–45_(S28D)-GST. In the absence of MX2, the capsid tubes exhibited well-defined edges with a clearly resolved continuous lattice (Fig. 3a). In contrast, the MX2_1–45_(S28D)-GST bound lattice displayed marked morphological changes: the tubes appeared heavily decorated and exhibited diduse or poorly defined edges and frequently clustered into irregular bundles (Fig. 3b).

**Figure 3:**
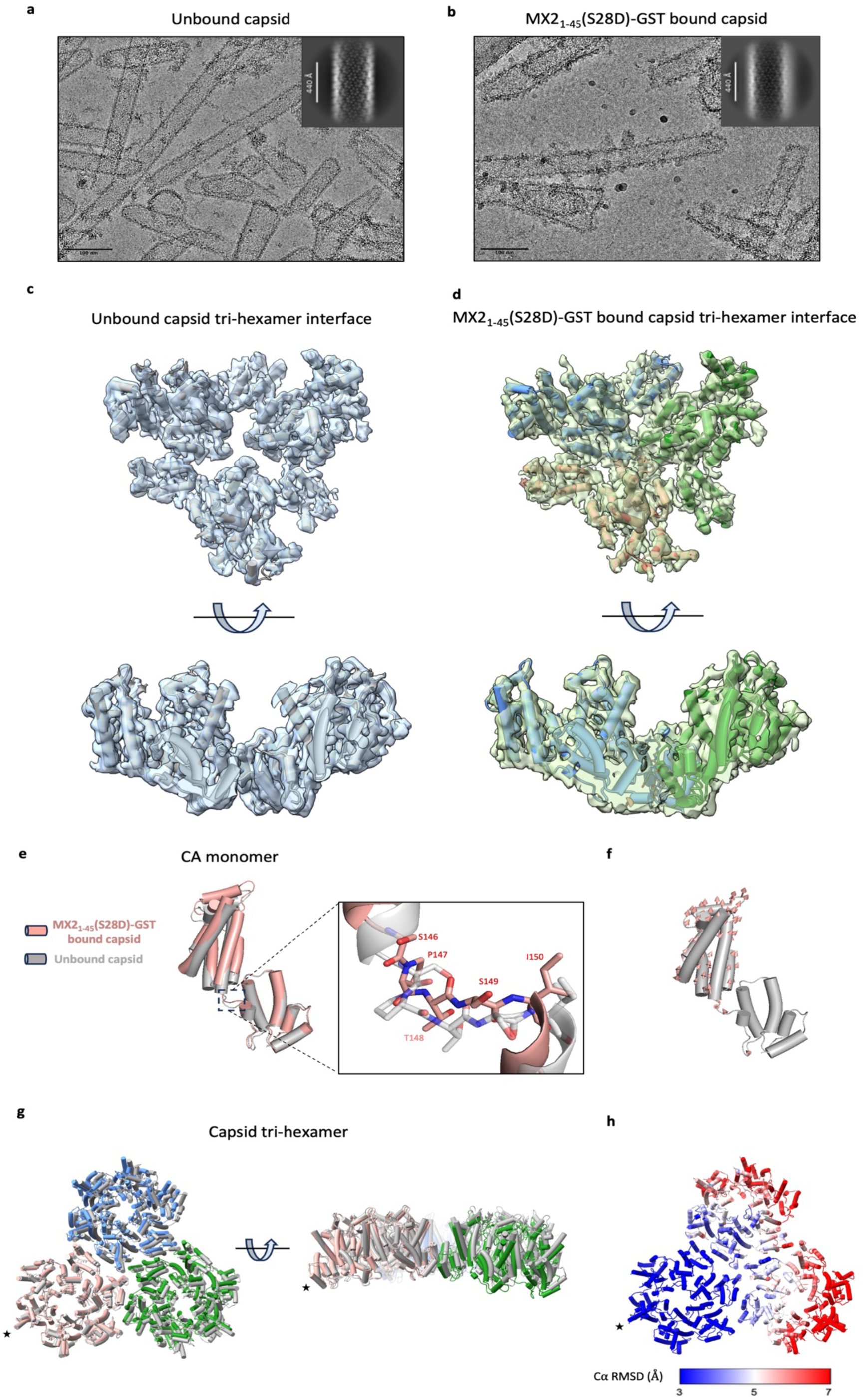
MX2 induces conformational changes on the capsid lattice. **a-b,** *In vitro* assembled capsid tubes in the absence (**a**) or presence (**b**) of MX2_1-45_(S28D)–GST, with representative 2D class averages shown. Images are representative of 9,131 (a) and 17,426 (**b**) micrographs, respectively. Scale bars, 100 nm. **c-d,** 3D cryo-EM reconstructions of the capsid tri-hexamer interface in the unbound (**c**) and MX2_1-45_(S28D)–GST bound (**d**) state with the refined atomic models. **e,** Atomic models of the unbound and MX2_1-45_(S28D)–GST bound capsid monomers aligned with residues 162-223. **f,** Model of the unbound capsid monomer with displacement vectors (salmon arrows) indicating movement to the MX2_1-45_(S28D)–GST bound model. **g,** Composite models of the unbound and MX2_1-45_(S28D)–GST bound capsid tri-hexamers aligned by the indicated CA monomer (*). Unbound capsid is coloured grey. MX2_1-45_(S28D)–GST capsid is coloured per hexamer. **h,** Unbound capsid tri-hexamer model coloured by per-residue Cα RMSD relative to the MX2_1-45_(S28D)–GST bound structure.

We observed distinct diderences in the helical structure of capsid tubes with and without MX2. Consistent with others^32,33^, 2D class averages of unbound capsid tubes displayed clear and distinct helical symmetries. By contrast, the 2D class averages of MX2_1–45_(S28D)-GST bound capsid tubes exhibited reduced lattice regularity and less defined helical symmetry (Fig. 3a-b, Extended Data Fig. 4), suggesting disruption to overall lattice organisation.

We next determined the structure of the capsid tubes, with and without MX2, by single-particle analysis (SPA). For each dataset, two maps were generated, centred on the hexamer with C6 symmetry applied; and on the tri-hexamer interface with no symmetry applied, respectively (See materials and methods for details). The hexamer-centred maps were determined to 3.33 Å resolution (unbound) and 3.71 Å resolution (MX2-bound); the tri-hexamer-centred maps were determined to 3.94 Å resolution (unbound) and 4.1 Å resolution (MX2-bound) (Extended Data Fig. 5-7, Extended Data Table 1).

To gain insights into long-range deformations of the capsid lattice upon MX2 binding, we employed the atomic models derived from the SPA reconstructions centred on the CA hexamer – they are at higher resolution and cover a larger region of the capsid lattice. Aligning the CTD of CA monomers from the unbound and MX2-bound capsid models reveals a conformational change in the relative orientation of the CA NTD and CA CTD upon MX2 binding (Fig. 3e-f), suggesting that MX2 binding induces structural rearrangements within the capsid lattice. Alignment of the unbound and MX2–bound capsid models revealed that the overall structure of the individual CA hexamers is very similar, with an average root mean square deviation (RMSD) of ∼1 Å (Extended Data Fig. 8). However, comparison of adjacent hexamers showed substantial change in their relative orientation, with RMSD values of 3–7 Å (Fig. 3g-h; Extended Data Movie 1). The greatest deviations were observed at the outer regions of the aligned model, indicating that MX2 binding perturbs the relative positioning of neighbouring hexamers. These results suggest that the molecular mechanism of MX2-mediated capsid destabilisation is underpinned by alterations at the hexamer-hexamer interface.

### MX2 residues 7–13 mediate binding at the tri-hexamer interface

In the hexamer-centred maps described above, density at the proposed MX2-binding site at the capsid tri-hexamer interface^24^ is diduse (likely due to the 6-fold symmetry applied), and did not permit atomic model building. To resolve this, we determined the maps of the CA tubes, unbound and bound to MX2_1–45_(S28D)-GST, centred on the tri-hexamer interface (See above), and without any symmetry applied. While the overall resolution of these maps was lower, clear density corresponding to MX2 was observed at the tri-hexamer interface in the MX2_1–45_(S28D)-GST map, not present in the map of unbound CA tubes (Fig. 3c-d).

An 8 amino acid peptide backbone could be modelled into this density, however the local resolution for this region was 4.5-6 Å (Extended Data Fig. 9b) such that the identity of the corresponding residues could not be determined from the density alone. To accommodate the MX2 sequence within the density, an 8–amino acid poly-alanine chain was then used as a template, and the sequence of the MX2 peptide (residues 1-45) was threaded into the model. Each of the 38 threaded models were subjected to 10 independent rounds of refinement, during which both the peptide and the three α-helices (helix 10 of CA) forming the pocket were allowed to move, and for each position the energy of the complex was calculated using Rosetta. This approach enabled evaluation of all potential registers of the MX2 1-45 sequence within the density and revealed one sequence (residues 7-PWPYRRRS-14) to have the lowest energy; accordingly, the density was consistent with placement of these residues (Extended Data Fig. 9a). MX2 residues upstream of P7 and downstream of S14 could not be resolved due to the lack of density. We also note that the density for the MX2 peptide is present at lower contour level than that of CA, that of CA, suggesting that MX2 was not occupying all tri-hexamer interfaces across the lattice in our dataset (Extended Data Fig. 9b).

In the resulting model, MX2 residues 7-14 are buried into the capsid tri-hexamer interface, with P7 positioned closest to the inside of the CA tube (Fig. 4a-b). Two distinct binding pockets are present in this structure: (1) MX2 residues R11 and R12 form salt bridges with residue E213 from adjacent CA monomers (Fig. 4c); and (2) MX2 residues P7, W8, P9, and Y10 form hydrophobic contacts with the three-helix bundle formed by CA helix 10 (Fig. 4d). As such, our data describe a new binding site for a ligand that engages the HIV-1 capsid lattice.

**Figure 4:**
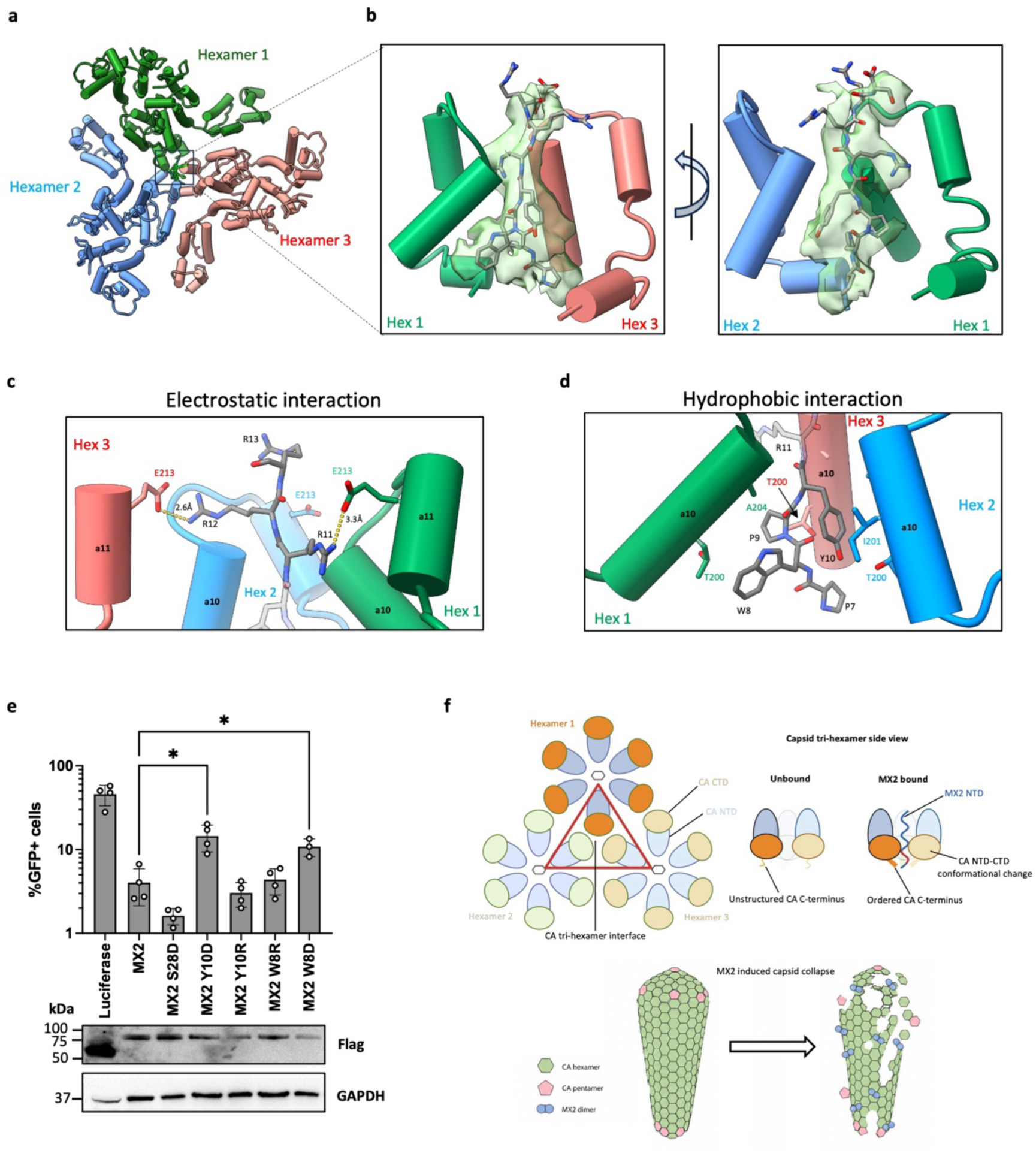
MX2 binds to the capsid tri-hexamer interface. **a,** Model of the MX2_1-45_(S28D)–GST bound capsid tri-hexamer interface. **b,** Model of MX2 residues 7-14 (grey) and CA residues 170-231 (green, blue and salmon) showing the cryo-EM density for MX2. **c-d,** Structure of the MX2 NTD electrostatic (**c**) and hydrophobic (**d**) interactions with the capsid tri-hexamer interface. **e,** U87-MG cells were transduced with EasiLV particles encoding FLAG-tagged luciferase, wild-type or mutant MX2 proteins. HIV-1/GFP infectivity was assessed 48 h post-infection by flow cytometry. Data represent mean ± SD from three independent biological replicates. Two-tailed paired *t*-test, * p<0.05. Protein expression was confirmed by immunoblot (below). **f,** Schematic of MX2 induced remodelling of the tri-hexamer interface (top) and MX2 induced capsid collapse (bottom).

Interestingly, additional density corresponding to the typically disordered CA CTD was observed in the MX2-bound map (Extended Data Fig. 10a). In the unbound capsid structure, no density is observed after residue G222, consistent with previously reported cryo-EM reconstructions^29,32,34,35^. In contrast, in the MX2-bound capsid map, density is clearly present for residues 223-231 (Extended Data Fig. 10b; Extended Data Movie 2). In our structure, MX2 residues P7 and W8 form hydrophobic contacts and stabilise these residues. Furthermore, inter-hexamer interactions between CA V230 and L231 from the C-terminal regions of adjacent hexamers (Extended Data Fig. 10c-d) are also present, supporting a model in which MX2 binding promotes the stabilisation of CA residues 223-231, therefore altering the inter-hexamer interface.

While the interaction between the RRR motif of MX2 and E213 (See above, Fig. 4c) had been proposed before, the hydrophobic interface (2) involving MX2 residues 7-10 (Fig. 4d) was unexpected. To assess the contribution of this interface to MX2 function, we introduced charge-altering substitutions at positions W8 and Y10. Incorporation of negatively charged aspartic acid (W8D and Y10D mutants) impaired HIV-1 restriction (Fig. 4e), highlighting the importance of these residues for antiviral activity. We note however that positively charged substitutions (W8R and Y10R mutants) had no significant phenotype (Fig. 4e). This could be attributable to the dominating impact of the electrostatic interface (1), to which such mutations could contribute, thus countering reduced interactions in the hydrophobic pocket.

## Discussion

Here, we define residues 1–45 of the MX2 NTD as the minimal region required for HIV-1 restriction. Single-particle cryo-EM analysis of the HIV-1 capsid bound to MX2_1–45_–GST revealed that electrostatic and hydrophobic contacts mediate MX2–capsid binding and stabilise the otherwise flexible CA C-terminus. MX2 binding markedly perturbs the organisation of the capsid lattice and triggers the collapse of CLPs and loss of lattice markers from virion-derived cores, suggesting that MX2 inhibits infection by binding to HIV-1 capsid cores in the cytoplasm and inducing premature fragmentation prior to nuclear import.

The MX2 NTD is predicted to be intrinsically disordered and its structure has remained unresolved in previous reports^16,17^. Consistent with this, only residues 7-14 of MX2 are resolved when bound to capsid, indicating that NTD residues downstream of the binding site do not strongly engage the lattice. Taken together with findings from mutagenesis of residues 35–45, these data suggest that downstream residues mainly serve as a flexible linker positioning residues 7-13 for capsid engagement. This agrees with previous reports that alanine scanning of the NTD outside the 11–13 RRR motif^19^ or partial NTD deletion (MX2_Δ26–50_)^36^ preserves antiviral activity. It is worth noting that phosphorylation at residue S28 enhances the antiviral activity of MX2. Despite using a phosphomimetic mutant (S28D) for our structure, this region of MX2 was not resolved. This may be attributable to this residue interacting with capsid upstream of the binding pockets described here, and the interaction not being observable in the averaging performed during SPA. Alternatively, this residue could have a yet-unidentified other role in MX2 restriction.

The positively charged 11–13 RRR motif is the principal determinant of capsid binding and HIV-1 restriction^19,20^. Neutralisation of this positive charge through W8D or Y10D substitutions (Fig. 4e) or phosphorylation/phosphomimetic mutation of residues S14, S17, and S18^22^ abolishes MX2 antiviral activity, underscoring the importance of electrostatic interactions with the negatively charged tri-hexamer interface. CA mutations at the tri-hexamer interface that alter this negatively charged interface (Extended Data Fig. 9c), such as G208R, T210K, or E212A/E213A, have been shown to hyperstabilise the lattice and impair virus infectivity as well as reduce MX2 binding^24,37^. Interestingly, the E212A/E213A mutation has further been shown to reduce core flexibility and impede nuclear import^37^, implicating the importance of this electrostatic environment for maintaining correct lattice stability. The 11–13 RRR motif engages E213, neutralising this negative potential and likely stabilising inter-hexamer interactions. In support of this, MX2 binding imparts order upon the otherwise flexible CA C-terminus^38^, establishing inter-hexamer contacts at the tri-hexamer interface (Extended Data Fig. 10). This MX2-induced remodelling of the C-terminus acts as a lever, that displaces adjacent hexamers and remodels the CA lattice.

The observation that MX2 induces disassembly of CLPs and virion-derived cores (Fig. 2) while capsid tubes remain intact (Extended Data Fig. 2) indicates that capsids with conical morphologies are preferentially susceptible to disruption. Our model of the MX2 NTD bound to capsid reveals that MX2 binding induces remodelling of the capsid lattice. The symmetric three-helix 10 bundle at the capsid tri-hexamer interface has been directly implicated in the variable curvature observed in native cores^35^, and we propose that MX2’s engagement with the capsid tri-hexamer interface restricts the conformational flexibility required to accommodate the full range of hexamer-hexamer curvatures observed within cores, and that this enforced rigidity ultimately results in the fracture and fragmentation of the capsid lattice.

The success of Lenacapavir as a therapeutic to treat and prevent HIV infection^39,40^ is a clear demonstration that capsid is an attractive target for antiviral compounds. Rational structure-based drug design could drastically accelerate drug development^41^, boosted recently by the emergence of Generative AI approaches^42^. Although the native capsid core is well characterised^43^, it may not fully capture the conformational landscape relevant for inhibitor design, which can involve alternative or inhibitor-stabilised states of CA that directly reveal conformations incompatible with the intact native structure. By obtaining the structure of core-incompatible, MX2-bound CA lattices, our work paves the way for the design of small-molecule inhibitors that bind to this interface in a similar way to MX2, and lead to capsid disassembly^44^.

Our model of MX2-mediated inhibition of HIV-1 infection provides a detailed molecular description of how a host restriction factor binds to, and disrupts, the HIV-1 capsid. This model permits to define the binding pocket to be targeted by novel capsid targeting inhibitors that mimic MX2 function.

## Materials and Methods

### Cells

HEK293T cells and U87-MG cells were obtained from the American Type Culture Collection (ATCC, CRL-1573 and HTB-14, respectively). All cell lines were cultured in Dulbecco’s modified Eagle medium (DMEM) supplemented with heat-inactivated foetal bovine serum (10%), L-glutamine, penicillin (100 U ml^−1^) and streptomycin (100 µg ml^−1^).

### Plasmids

pEasiLV containing luciferase, wild-type MX2, MX2 mutants and Fv1b(NTD_MX2_) chimeras, were produced by standard cloning or site directed mutagenesis. CypA-DsRed and VPR-IN-mNeonGreen plasmids were gifts from the Francis Laboratory (Florida State University).

### Viral production

EasiLV particles were produced by transfection with pCMV-ΔR8.91 (p8.91), pEasiLV, pEasiLV, pptTRKrab and pMD.G at a ratio of 1:1:0.5:0.25, respectively, with TransIT-VirusGEN (Mirus) reagent at a 2:1 ratio with DNA. Cell culture medium was replaced after 24 hours and after a further 24 hours the media filtered through 0.45 µm cut-od filter and used to directly transduce U87-MG cells. After 6 hours, the media was replaced and doxycycline added (0.5 µg ml^−1^, Sigma-Aldrich) to induce transgene expression. U87-MG cells were challenged with a lenti-GFP vectors and infectivity measured by flow cytometry (NovoCyte) after 48 hours.

HIV-1 virions were produced by following the same protocol used by Wei et al (2022)^33^.

HIV-1 pseudoparticles incorporating CypA-DsRed and VPR-IN-mNeonGreen were produced by transfection with pR8_Δ_Env, pcREV, CypA-DsRed, VPR-IN-mNeonGreen at a ratio of 1:1:0.5:0.25, respectively, with TransIT-VirusGEN (Mirus) reagent at a 2:1 ratio with DNA. After 48 hours, media was collected and filtered through a 0.45 µM syringe filter (Sartorius) before being concentrated by centrifugation.

### Fluorescence assay

HIV-1 pseudoparticles were allowed to bind to an 8-well chambered coverslide. Particles were imaged for in 2 minutes intervals for 40 minutes at room temperature. Following the first acquisition, 50 µL PBS containing MX2 proteins, IP6, and saponin was added to give final concentrations of 10 µM, 20 µM, and 50 µg mL^−1^, respectively. Particles were analysed using Fiji by applying Maximum Entropy thresholding.

### Immunoblotting

U87-MG cells were lysed in buder containing 10 mM Tris-HCl pH 8, 150 mM NaCl, 1 mM EDTA, 1% Triton X-100 and 0.1% sodium deoxycholate. Lysates were briefly sonicated, boiled for 10 min and resolved by SDS-PAGE before being analysed by immunoblotting using primary antibodies specific to Flag (mouse monoclonal M2, Sigma-Aldrich) followed by secondary anti-mouse horseradish peroxidase-conjugated antibody.

### Protein expression and purification

HIV-1 CA was expressed and purified as described^45^. MX2-GST proteins were expressed within Escherichia coli BL21 and purified by Ni-NTA adinity and size exclusion chromatography (Superdex 200 pg, Cytiva).

### Capsid pulldown

Pseudotyped HIV-1 was permeabilised with 0.5% Triton X-100 in STE buder (10 mM Tris, pH 7.4, 100 mM NaCl, 1 mM EDTA) and incubated with MX2(1–45)–GST or MX2_1–45_(RRR/AAA)-GST (0.05–2.5 µM) at 4 °C. Complexes were captured using Glutathione Sepharose 4B beads (GE healthcare), washed, and eluted by boiling in STE buder containing 0.1% SDS. Bound capsid was quantified by p24 ELISA (Zeptometrix) following the manufacturer’s instructions.

### CA assembly and MX2 binding

CA was assembled into tubular capsid oligomers by incubating the protein (21 mg mL^−1^) in 5 mM IP6, 50 mM Tris pH 8.0, 150 mM NaCl, 5 mM <u>f3</u>-mercaptoethanol (BME) at 37°C for 1-2 hours. Assembled tubular capsid particles were centrifuged for 10 minutes at 15,000 RPM and resuspended in 200 µM IP6, 50 mM Tris pH 8.0, 150 mM NaCl, 5 mM BME. MX2 (7 mg mL^−1^) or buder was added to pre-assembled capsid particles and incubated for 1 hour at 4°C.

CLPs were assembled by incubating the protein (21 mg mL^−1^) in 500 µM IP6, 50 mM MES pH 6.0, 150 mM NaCl at 37°C overnight. CLPs were incubated with MX2_1–45_GST or MX2_1–45_(RRR/AAA)-GST at 37°C for 1 hour.

### Negative-stain EM

Samples (3 µL) were applied to glow discharged 300 mesh copper grids with a carbon film (electron Microscopy Sciences) and incubated for 30 seconds. Grids were washed in a 30 µL drop of 2% uranyl acetate (UA) and blotted briefly. Grids were then placed on a second drop of UA and incubated for 2 minutes before being blotted dry. Images were collected at on a Tecnai T12 Spirit TEM operating at 120 kV, equipped with a 2 K Eagle camera.

### Single-particle cryo-EM sample preparation and data collection

Tubular capsid oligomers with and without MX2 were diluted 10-fold with 100 µM IP6, 50 mM Tris pH 8.0, 150 mM NaCl, 5 mM BME and 3 µL immediately applied to glow-discharged holey carbon grids (Quantifoil R2/2, 300-mesh). Grids were blotted and plunge frozen in liquid ethane using a Leica GP1 freeze-plunger.

Cryo-EM data was collected using a Krios G3i microscope (ThermoFisher) operating at 300 kV and equipped with a K3 direct electron detector (Gatan). Data was a collected using EPU (ThermoFisher) using a pixel size of 1.078 Å in super resolution mode. For MX2_1–45_(S28D)-GST bound capsid a total of 17,426 movies were collected with a total dose of 45 electrons/Å^2^. For unbound capsid a total of 9131 movies were collected with a total dose of 50 electrons/Å^2^. All movies were collected over 40 frames with a defocus range of -0.7 to -2.2 µm.

### Image processing

For all movies, motion correction and CTF estimation were performed in cryoSPARC^46^ v4.5 (Structura Biotechnology) using the patch motion correction and patch CTF estimation respectively. Initial particles were picked manually to generate references for subsequent template-based picking.

For unbound and MX2_1–45_(S28D)-GST bound capsid datasets, particles were extracted with a box size of 252 pixels (227 Å). Sequential rounds of 2D classification were performed to remove junk particles, after which selected particles were re-extracted with a box size of 400 pixels (440 Å). For the MX2_1–45_(S28D)-GST bound capsid, a single round of heterogenous refinement with C1 symmetry was performed to remove additional junk particles. Homogenous refinement and non-uniform refinement were performed with C6 symmetry using an initial input model of 7 capsid hexamers (PDB 3J34).

For the maps centred at the capsid tri-hexamer interface, particles were re-aligned on the tri-hexamer interface and re-extracted with a box size of 252 pixels (227 Å). Homogenous refinement and non-uniform refinement were performed with C1 symmetry. For the maps centred at the capsid tri-hexamer interface, particles were re-aligned on the tri-hexamer interface and re-extracted with a box size of 252 pixels (227 Å). Homogenous refinement and non-uniform refinement were performed with C1 symmetry.

### Model building, refinement and peptide modelling

Initial model of a CA hexamer was generated using AlphaFold3^47^ and refined into the density using ISOLDE^48^ in UCSF ChimeraX^49^, followed by Real Space Refinement in Phenix^50^.

To model the MX2 sequence bound to CA, thirty-nine candidate MX2 sequence registrations were threaded onto an 8-residue polyalanine backbone built into the resolved density and subjected to real-space refinement in Rosetta^51^. Refinement used the default Rosetta energy function together with a density-fit term. Peptide atoms and CA residues within 10 Å of the peptide were allowed to move, while the remainder of the structure was fixed. For each registration, 10 refined models were generated, yielding 390 models in total. The best-scoring registration showed both minimum and mean energies

>7 kcal mol⁻¹ lower than the second-best registration.

## ACKNOWLEDGMENTS

We thank Benjamin Chen, Zandrea Ambrose, Ashwanth Francis, Shamar Lale and Stelios Papaioannou for the provision of reagents. The work was funded by the Wellcome Trust (106223/Z/14/Z and 222433/Z/21/Z to M.H.M.), the Medical Research Council (MR/S023747/1 to M.H.M and L.A.) and the National Institutes of Health (U54 AI150472 and U54 AI170855 to M.H.M and M.K., R01 AI157802 to M.K.), J.R.C.B acknowledges funding from the Human Frontier Science Program (RGY0080/2021). A.G. was supported by a Medical Research Council – King’s College London Doctoral Training Partnership in Biomedical Sciences (MR/W006820/1), N.A. was funded by the Wellcome Trust PhD programme in Cell Therapies and Regenerative Medicine (108874/Z/15/Z). We thank the Nikon Centre at King’s College London for their expertise and guidance in utilizing high-resolution confocal microscopy. We thank members of the Bergeron, Malim, Apolonia and Atherton laboratories for helpful discussions. Cryo-EM data was collected at the LonCEM facility, where we thank Nora Cronin for support.

## Data availability

The coordinates and EM maps of the unbound and MX2_1-45_(S28D)-GST bound CA have been deposited in the PDB and EMDB databases with the following accession codes: the unbound CA hexamer, PDB: 30OE and EMDB: EMD-57889; MX2_1-45_(S28D)-GST bound CA hexamer, PDB: 30OD and EMDB: EMD-57888; unbound CA tri-hexamer interface PDB: 30OG and EMDB: EMD-57891; and MX2_1-45_(S28D)-GST bound CA tri-hexamer interface, PDB: 30OF and EMDB: EMD-57890. Source data are provided with this paper.

## References

1. Rasaiyaah, J. et al. HIV-1 evades innate immune recognition through specific cofactor recruitment. Nature 503, 402–405 (2013).

2. Lahaye, X. et al. The capsids of HIV-1 and HIV-2 determine immune detection of the viral cDNA by the innate sensor cGAS in dendritic cells. Immunity 39, 1132–1142 (2013).

3. Ganser, B. K., Li, S., Klishko, V. Y., Finch, J. T. & Sundquist, W. I. Assembly and Analysis of Conical Models for the HIV-1 Core. Science 283, 80–83 (1999).

4. Liu, C. et al. Cyclophilin A stabilizes the HIV-1 capsid through a novel non-canonical binding site. Nat. Commun. 7, 10714 (2016).

5. Fu, L. et al. HIV-1 capsids enter the FG phase of nuclear pores like a transport receptor. Nature 626, 843–851 (2024).

6. Dickson, C. F. et al. The HIV capsid mimics karyopherin engagement of FG-nucleoporins. Nature 626, 836–842 (2024).

7. Kreysing, J. P. et al. Passage of the HIV capsid cracks the nuclear pore. Cell 188, 930–943.e21 (2025).

8. Achuthan, V. et al. Capsid-CPSF6 interaction licenses nuclear HIV-1 trafficking to sites of viral DNA integration. Cell Host Microbe 24, 392–404.e8 (2018).

9. Goujon, C. et al. Human MX2 is an interferon-induced post-entry inhibitor of HIV-1 infection. Nature 502, 559–562 (2013).

10. Kane, M. et al. Mx2 is an interferon induced inhibitor of HIV-1 infection. Nature 502, 563–566 (2013).

11. Crameri, M. et al. MxB is an interferon-induced restriction factor of human herpesviruses. Nat. Commun. 9, 1980 (2018).

12. Wang, Y.-X. et al. Interferon-inducible MX2 is a host restriction factor of hepatitis B virus replication. J. Hepatol. 72, 865–876 (2020).

13. Yi, D.-R. et al. Human MxB Inhibits the Replication of Hepatitis C Virus. J. Virol. 93, 10.1128/jvi.01285-18 (2018).

14. Ramachandran, R. & Schmid, S. L. The dynamin superfamily. Curr. Biol. 28, R411–R416 (2018).

15. Ferguson, S. M. & De Camilli, P. Dynamin, a membrane remodelling GTPase. Nat. Rev. Mol. Cell Biol. 13, 75–88 (2012).

16. Fribourgh, J. L. et al. Structural Insight into HIV-1 Restriction by MxB. Cell Host Microbe 16, 627 (2014).

17. Alvarez, F. J. D. et al. CryoEM structure of MxB reveals a novel oligomerization interface critical for HIV restriction. Sci. Adv. 3, e1701264 (2017).

18. Serrero, M. C. et al. The interferon-inducible GTPase MxB promotes capsid disassembly and genome release of herpesviruses. eLife 11, e76804 (2022).

19. Goujon, C., Greenbury, R. A., Papaioannou, S., Doyle, T. & Malim, M. H. A Triple-Arginine Motif in the Amino-Terminal Domain and Oligomerization Are Required for HIV-1 Inhibition by Human MX2. J. Virol. 10.1128/jvi.00169-15 (2015) doi:10.1128/jvi.00169-15.

20. Betancor, G. et al. The GTPase Domain of MX2 Interacts with the HIV-1 Capsid, Enabling Its Short Isoform to Moderate Antiviral Restriction. Cell Rep. 29, 1923–1933.e3 (2019).

21. Dicks, M. D. J. et al. Multiple components of the nuclear pore complex interact with the amino-terminus of MX2 to facilitate HIV-1 restriction. PLOS Pathog. 14, e1007408 (2018).

22. Betancor, G. et al. MX2-mediated innate immunity against HIV-1 is regulated by serine phosphorylation. Nat. Microbiol. 6, 1031–1042 (2021).

23. Betancor, G. et al. MX2 Viral Substrate Breadth and Inhibitory Activity Are Regulated by Protein Phosphorylation. mBio 13, e01714–22.

24. Smaga, S. S. et al.. MxB Restricts HIV-1 by Targeting the Tri-hexamer Interface of the Viral Capsid. Struct. Lond. Engl. 1993 27, 1234–1245.e5 (2019).

25. Busnadiego, I. et al. Host and Viral Determinants of Mx2 Antiretroviral Activity. J. Virol. 88, 7738–7752 (2014).

26. Mallery, D. L. et al. IP6 is an HIV pocket factor that prevents capsid collapse and promotes DNA synthesis. eLife 7, e35335.

27. Renner, N. et al. A lysine ring in HIV capsid pores coordinates IP6 to drive mature capsid assembly. PLoS Pathog. 17, e1009164 (2021).

28. Dick, R. A. et al. Inositol phosphates are assembly co-factors for HIV-1. Nature 560, 509–512 (2018).

29. Schirra, R. T. et al. A molecular switch modulates assembly and host factor binding of the HIV-1 capsid. Nat. Struct. Mol. Biol. 30, 383–390 (2023).

30. Francis, A. C., Marin, M., Shi, J., Aiken, C. & Melikyan, G. B. Time-Resolved Imaging of Single HIV-1 Uncoating In Vitro and in Living Cells. PLoS Pathog. 12, e1005709 (2016).

31. Zhao, G. et al. Mature HIV-1 capsid structure by cryo-electron microscopy and all-atom molecular dynamics. Nature 497, 643–646 (2013).

32. Wei, G. et al. Prion-like low complexity regions enable avid virus-host interactions during HIV-1 infection. Nat. Commun. 13, 5879 (2022).

33. Zhu, Y. et al. Structural basis for HIV-1 capsid adaption to rescue IP6-packaging deficiency. bioRxiv 2025.02.09.637297 (2025) doi:10.1101/2025.02.09.637297.

34. Stacey, J. C. V. et al. Two structural switches in HIV-1 capsid regulate capsid curvature and host factor binding. Proc. Natl. Acad. Sci. U. S. A. 120, e2220557120.

35. Moschonas, G. D. et al. MX2 forms nucleoporin-comprising cytoplasmic biomolecular condensates that lure viral capsids. Cell Host Microbe 32, 1705–1724.e14 (2024).

36. Deshpande, A. et al. Elasticity of the HIV-1 core facilitates nuclear entry and infection. PLOS Pathog. 20, e1012537 (2024).

37. Zhu, Y. et al. Structural basis for HIV-1 capsid adaption to a deficiency in IP6 packaging. Nat. Commun. 16, 8152 (2025).

38. Balter, M. New Hope in HIV Disease. Science 274, 1988–1991 (1996).

39. De Clercq, E. et al. Lenacapavir: A capsid inhibitor for HIV-1 treatment and prevention. Biochem. Pharmacol. 240, 117125 (2025).

40. Nero, T. L., Parker, M. W. & Morton, C. J. Protein structure and computational drug discovery. Biochem. Soc. Trans. 46, 1367–1379 (2018).

41. Zhong, Z. & Durrant, J. D. Generative AI in structure-based drug discovery. BMC Biol. 10.1186/s12915-026-02575-x (2026) doi:10.1186/s12915-026-02575-x.

42. Morling, K. L., ElGhazaly, M., Milne, R. S. B. & Towers, G. J. HIV capsids: orchestrators of innate immune evasion, pathogenesis and pandemicity. J. Gen. Virol. 106, 002057 (2025).

43. Zhang, C., Li, B., Li, J., Zhang, H. & Wu, Y. New Advances in Anti-HIV-1 Strategies Targeting the Assembly and Stability of Capsid Protein. Int. J. Mol. Sci. 26, 5819 (2025).

44. Hung, M. et al. Large-Scale Functional Purification of Recombinant HIV-1 Capsid. PLOS ONE 8, e58035 (2013).

45. Punjani, A., Rubinstein, J. L., Fleet, D. J. & Brubaker, M. A. cryoSPARC: algorithms for rapid unsupervised cryo-EM structure determination. Nat. Methods 14, 290–296 (2017).

46. Abramson, J. et al. Accurate structure prediction of biomolecular interactions with AlphaFold 3. Nature 630, 493–500 (2024).

47. Croll, T. I. ISOLDE: a physically realistic environment for model building into low-resolution electron-density maps. Acta Crystallogr. Sect. Struct. Biol. 74, 519–530 (2018).

48. Pettersen, E. F. et al. UCSF ChimeraX: Structure visualization for researchers, educators, and developers. Protein Sci. Publ. Protein Soc. 30, 70–82 (2021).

49. Afonine, P. V. et al. New tools for the analysis and validation of cryo-EM maps and atomic models. Acta Crystallogr. Sect. Struct. Biol. 74, 814–840 (2018).

50. Wang, R. Y.-R. et al. Automated structure refinement of macromolecular assemblies from cryo-EM maps using Rosetta. eLife 5, e17219 (2016).

