## Extended Data Figure for "MX2 Mediates Collapse of the HIV-1 Capsid"

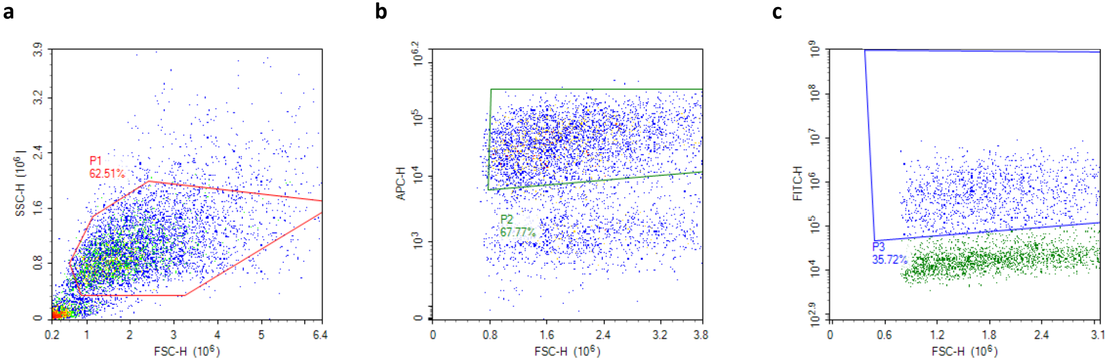


**Extended Data Figure 1:**

**a,** Live cells were first gated by forward scatter (FSC-H) and side scatter (SSC-H) to exclude debris and select intact populations. **b,** E2-crimson–positive cells were then identified using APC fluorescence (APC-H). **c,** GFP-positive cells were subsequently gated based on FITC fluorescence (FITC-H).

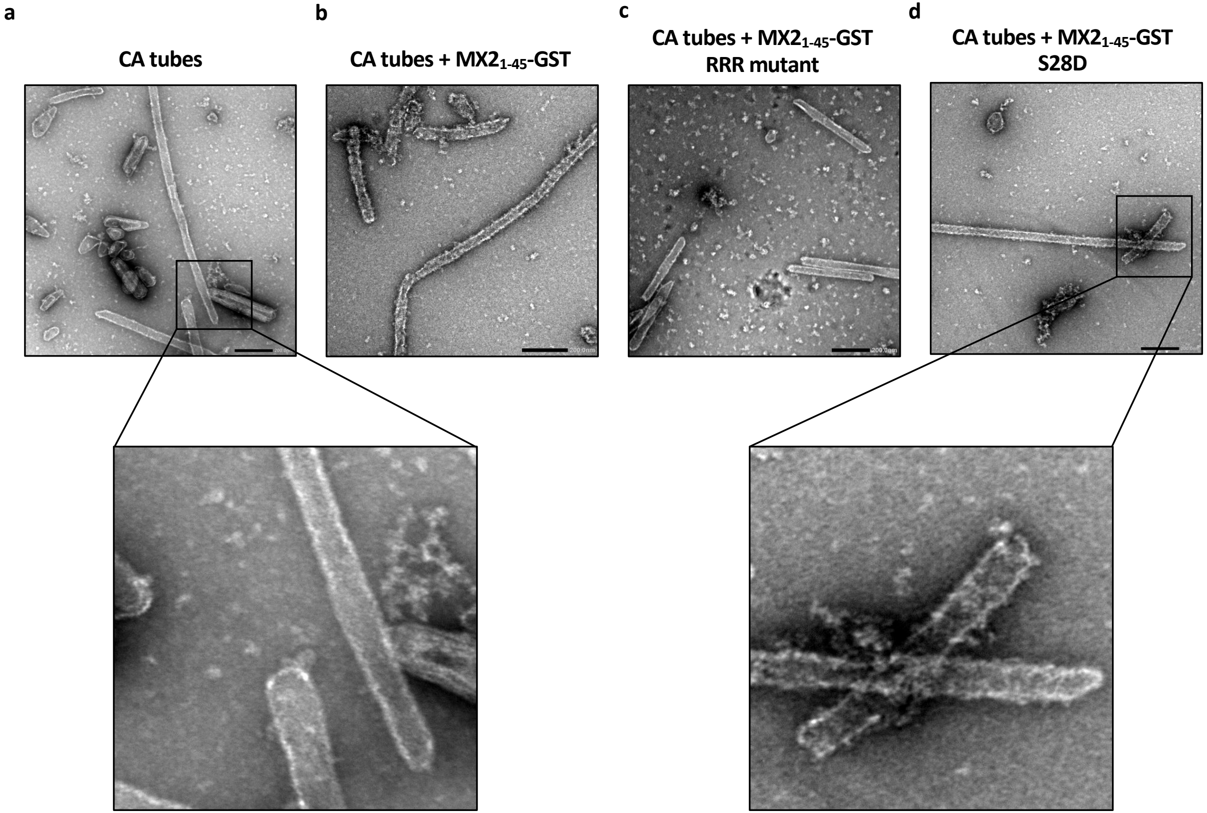


**Extended Data Figure 2: Morphology of MX2 bound capsid tubes**

**a-d,** Negative-stain TEM micrographs of capsid tubes assembled from CA in 5 mM IP6, 50 mM Tris–HCl (pH 8.0) and 150 mM NaCl. Tubes were pelleted by centrifugation, resuspended in buffer containing 200 µM IP6, and incubated with the indicated MX2(1–45)–GST constructs. Scale bars, 200 nm.

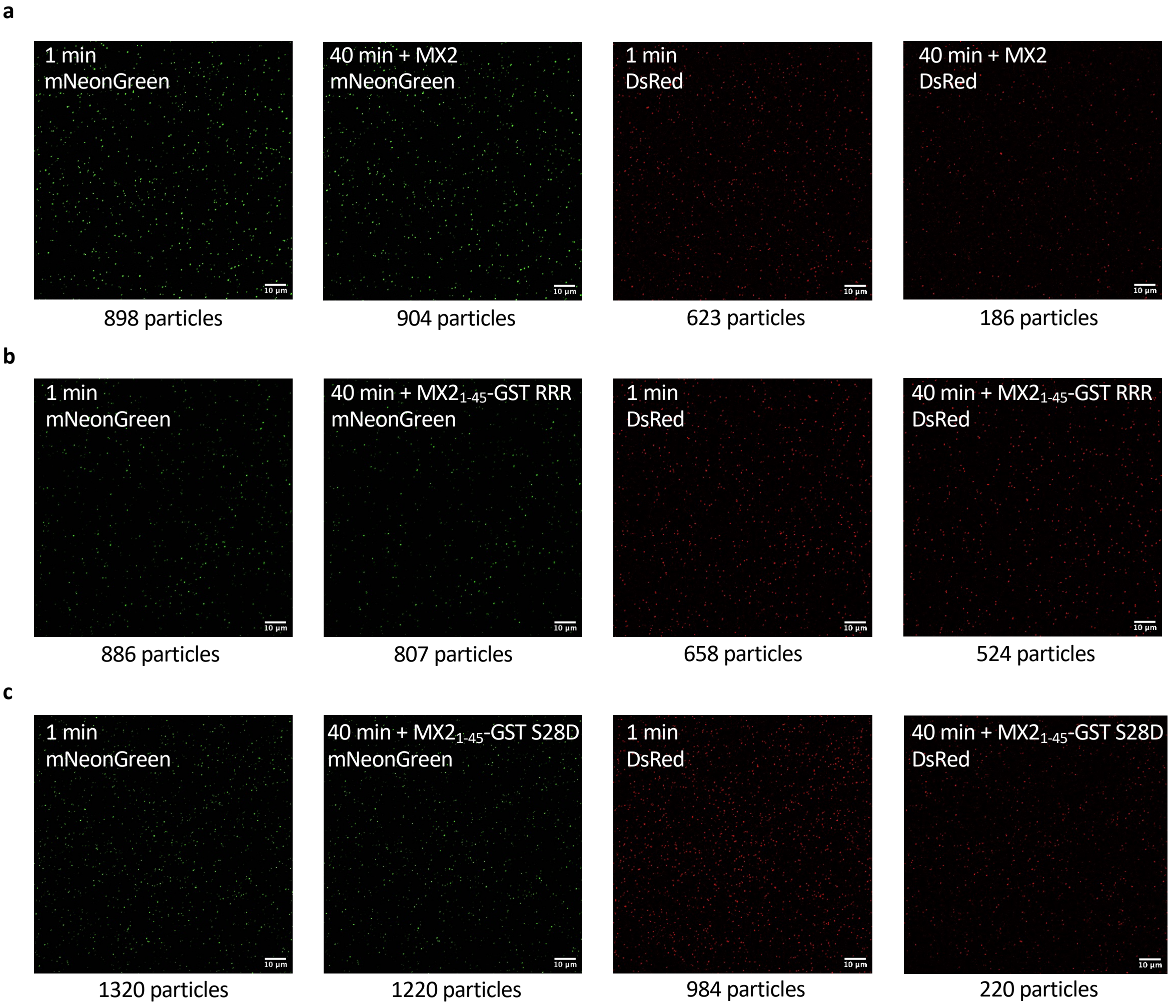


**Extended Data Figure 3: MX2 causes loss of capsid lattice marker**

**a-c,** Virus-like particles containing cores co-labelled with IN-mNeonGreen and CypA-DsRed. Bond particles were incubated with permeabilisation buffer containing MX2$_{1-45}$-GST (**a**), MX2$_{1-45}$-GST 11-13AAA (**b**), or MX2$_{1-45}$-GST S28D (**c**) for 40 minutes.

**a**

**Unbound capsid**

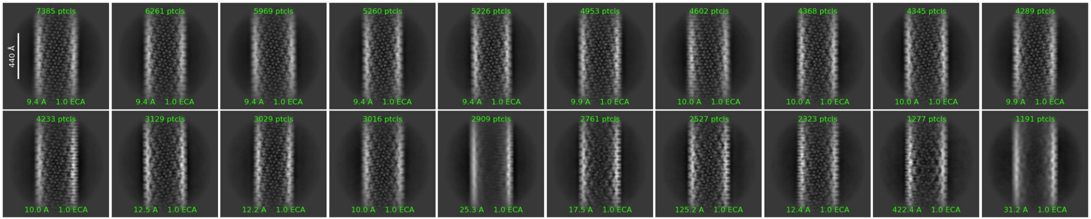


**b**

**MX2$_{1\text{-}45}$-GST S28D bound capsid**

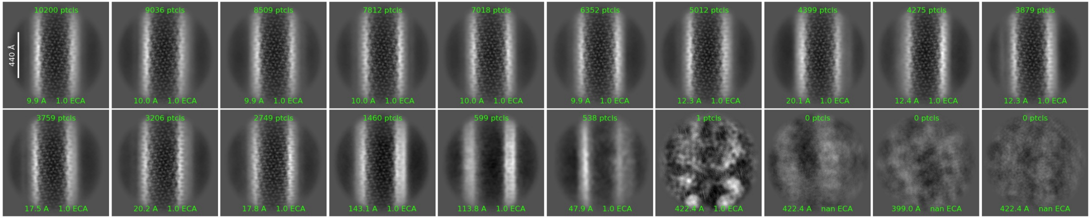


**Extended Data Figure 4: MX2 perturbs the helical structure of CA tubes**

**a-b,** Representative 2D classes from ~70,000 particles extracted with a box size of 960 px (1056 Å) for unbound (**a**) and MX2$_{1\text{-}45}$-GST S28D bound (**b**) capsid.

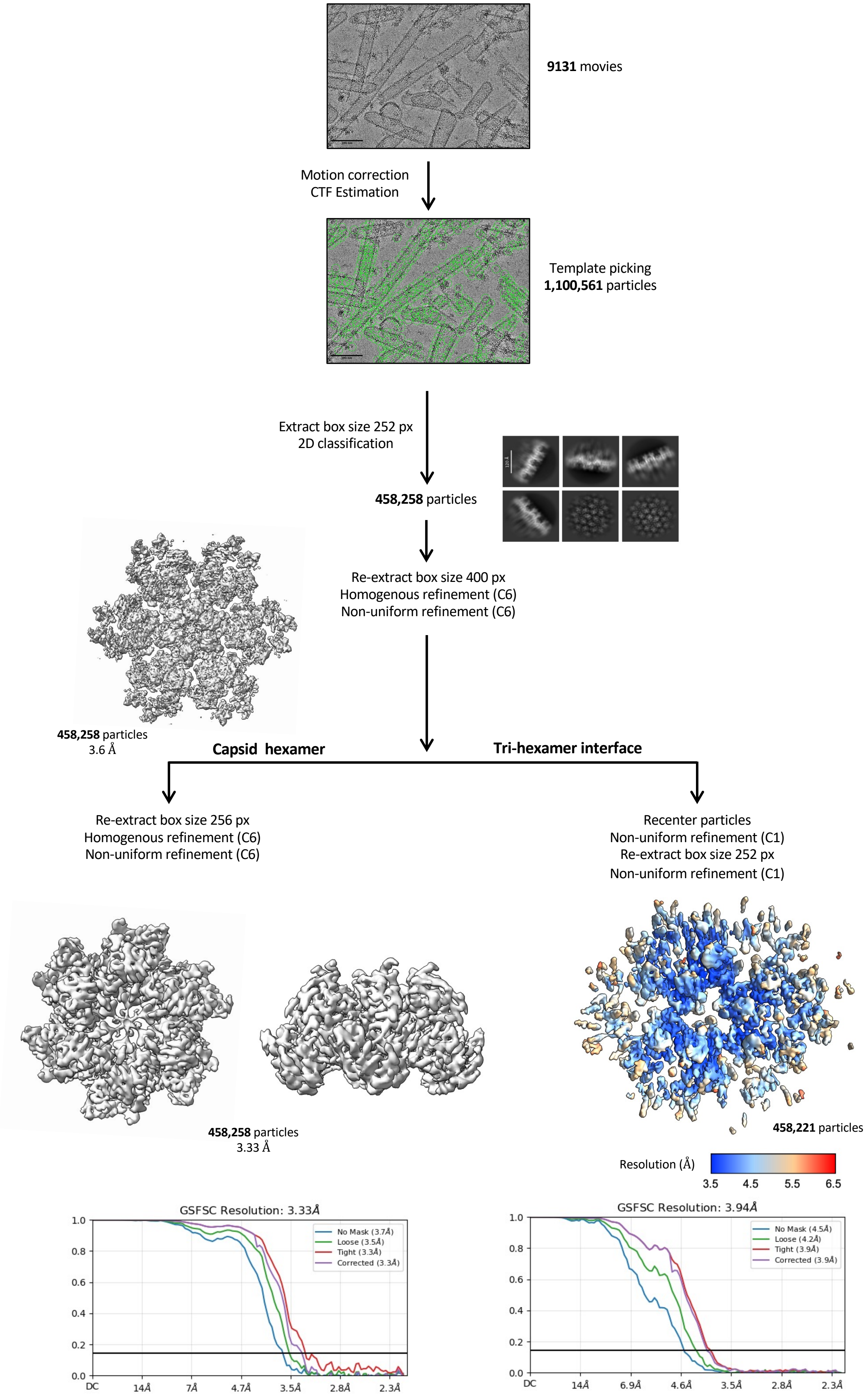


**Extended Data Figure 5**: **Cryo-EM processing pipeline of unbound capsid reconstructions**

Flowchart of the SPA processing pipeline for the unbound capsid reconstructions. Template-based picking resulted in 1,100,561 particles from 9131 micrographs. 2D classification performed to remove junk particles before 458,258 particles were used to generate an initial model of 7 CA hexamers centred on a CA hexamer, obtaining a 3.6 Å resolution map. These particles were used to refine the maps of the CA hexamer (C6 symmetry applied) and the tri-hexamer interface (no symmetry applied) achieving a global resolutions of 3.33 Å and 3.94 Å, respectively. The map of the tri-hexamer interface has been coloured by local resolution.

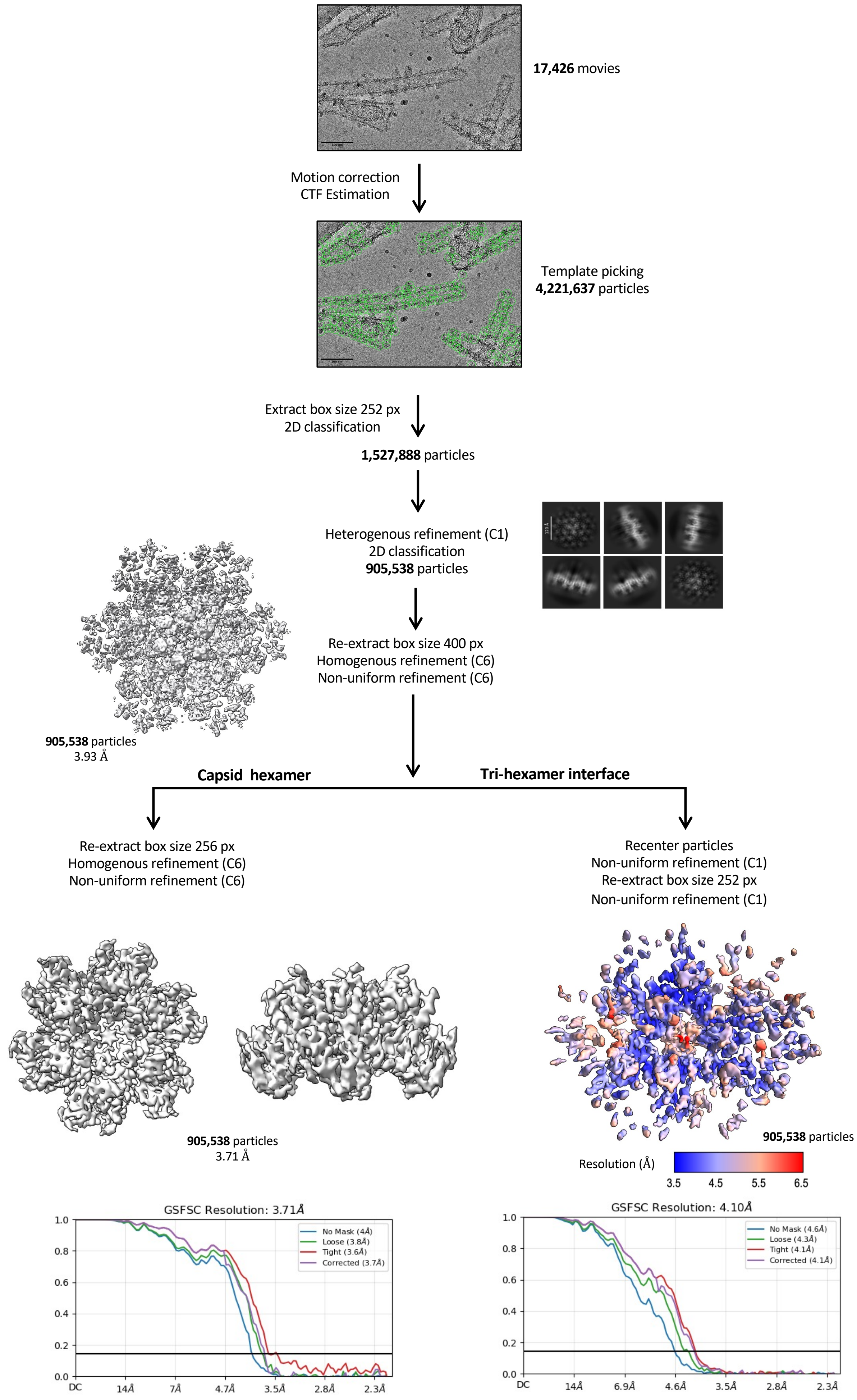


**Extended data figure 6: Cryo-EM processing pipeline of MX2 bound capsid reconstructions**

Flowchart of the processing pipeline for the $MX2_{1\text{-}45}$-GST S28D bound capsid reconstructions. Template-based picking resulted in 4,211,637 particles from 17,426 micrographs. 2D classification and heterogenous refinement were performed to remove junk particles. A total of 905,538 particles were used to generate an initial model of 7 CA hexamers centred on a CA hexamer, obtaining a 3.93 Å resolution map. These particles were used to refine the maps of the CA hexamer (C6 symmetry applied) and the tri-hexamer interface (no symmetry applied) achieving a global resolutions of 3.71 Å and 4.1 Å, respectively. The map of the tri-hexamer interface has been coloured by local resolution.

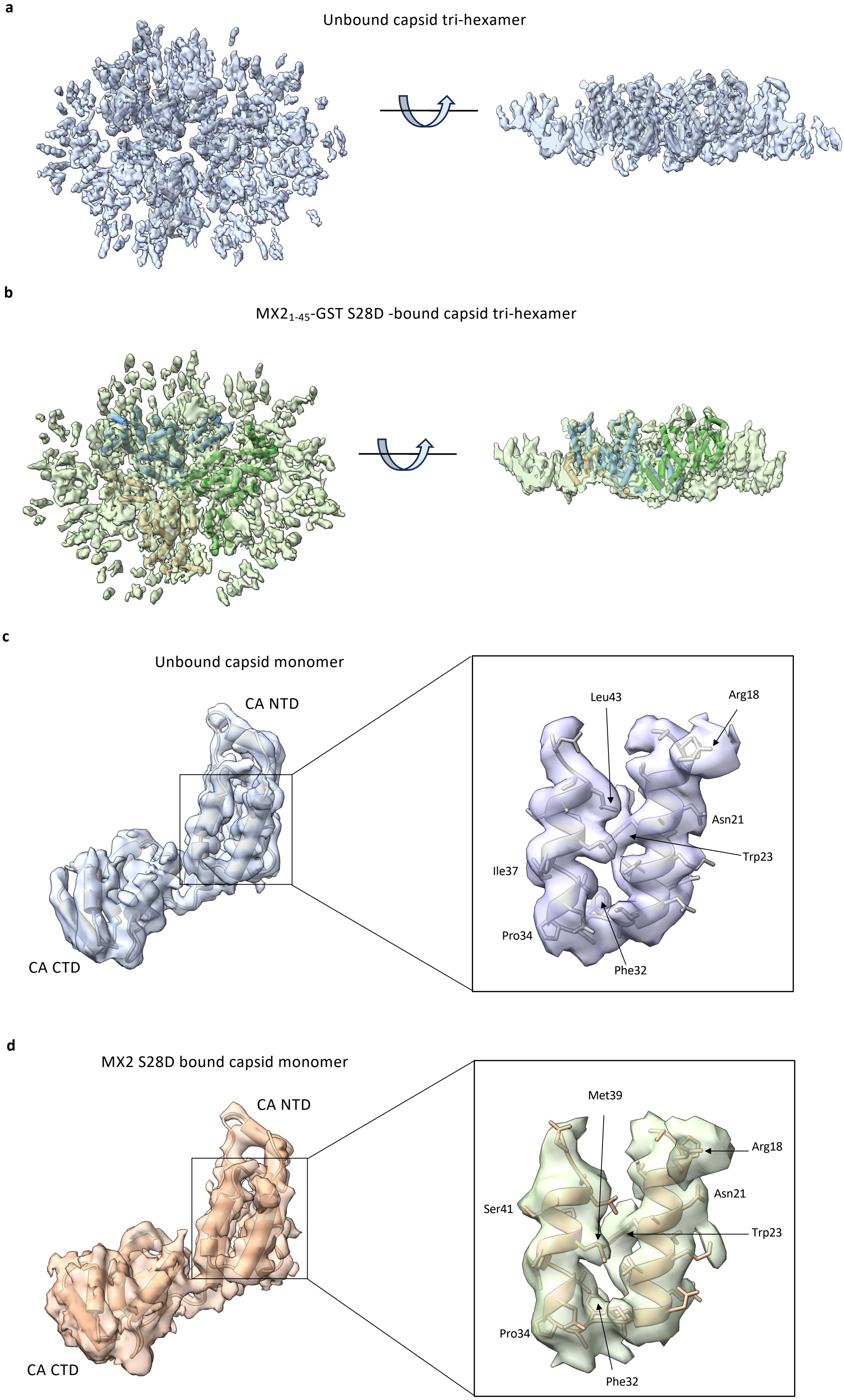


**Extended Data Figure 7**: **Structure of the unbound and MX2 bound tri-hexamer interface**
**a-b,** 3D cryo-EM reconstructions of the capsid tri-hexamer in the unbound (**a**) and MX2$_{1\text{-}45}$-GST S28D bound (**b**) state with the refined atomic models of the tri-hexamer interface. **c-d,** Atomic models of the unbound (**c**) and MX2$_{1\text{-}45}$-GST S28D–bound (**d**) CA monomer extracted from the capsid tri-hexamer models in **a–b**. The cryo-EM density maps are the same as in **a–b**, zoned to highlight the the modelled CA monomers at the tri-hexamer interface.

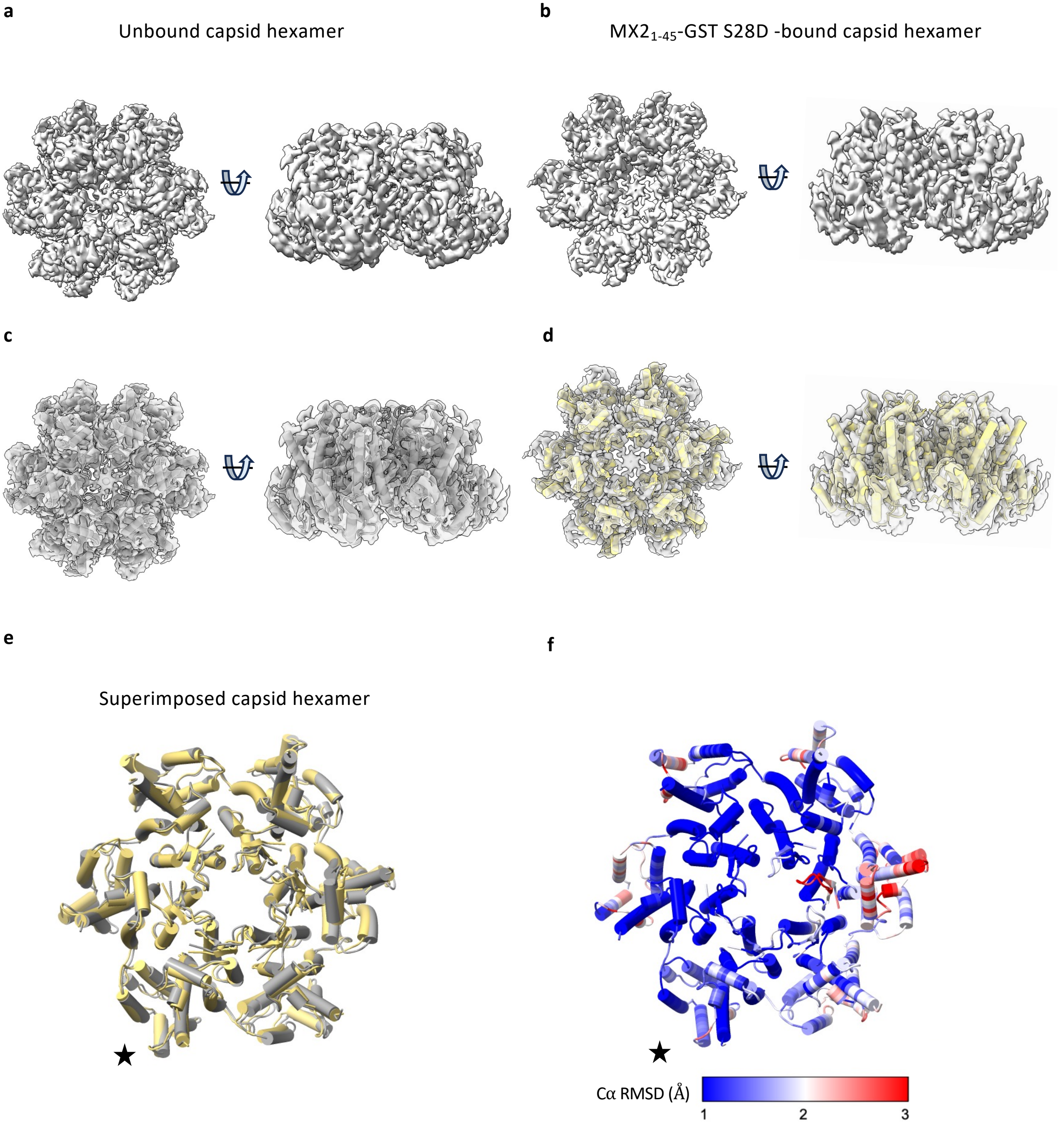


**Extended Data Figure 8: MX2 binding does not alter the structure of the CA hexamer**

**a-b,** Cryo-EM density maps of the unbound (**a**) and MX2$_{1-45}$-GST S28D bound (**b**) capsid hexamers. **c–d,** Same maps as in **a,** and **b,** with fitted atomic models for the unbound (grey, **c**) and MX2$_{1-45}$-GST S28D bound (yellow, **d**) capsid hexamers. **e,** Models of the unbound and MX2$_{1-45}$-GST S28D bound capsid hexamers aligned by the indicated CA monomer (*). **f,** Unbound capsid hexamer model coloured by per-residue Cα RMSD relative to the MX2$_{1-45}$-GST S28D bound structure.

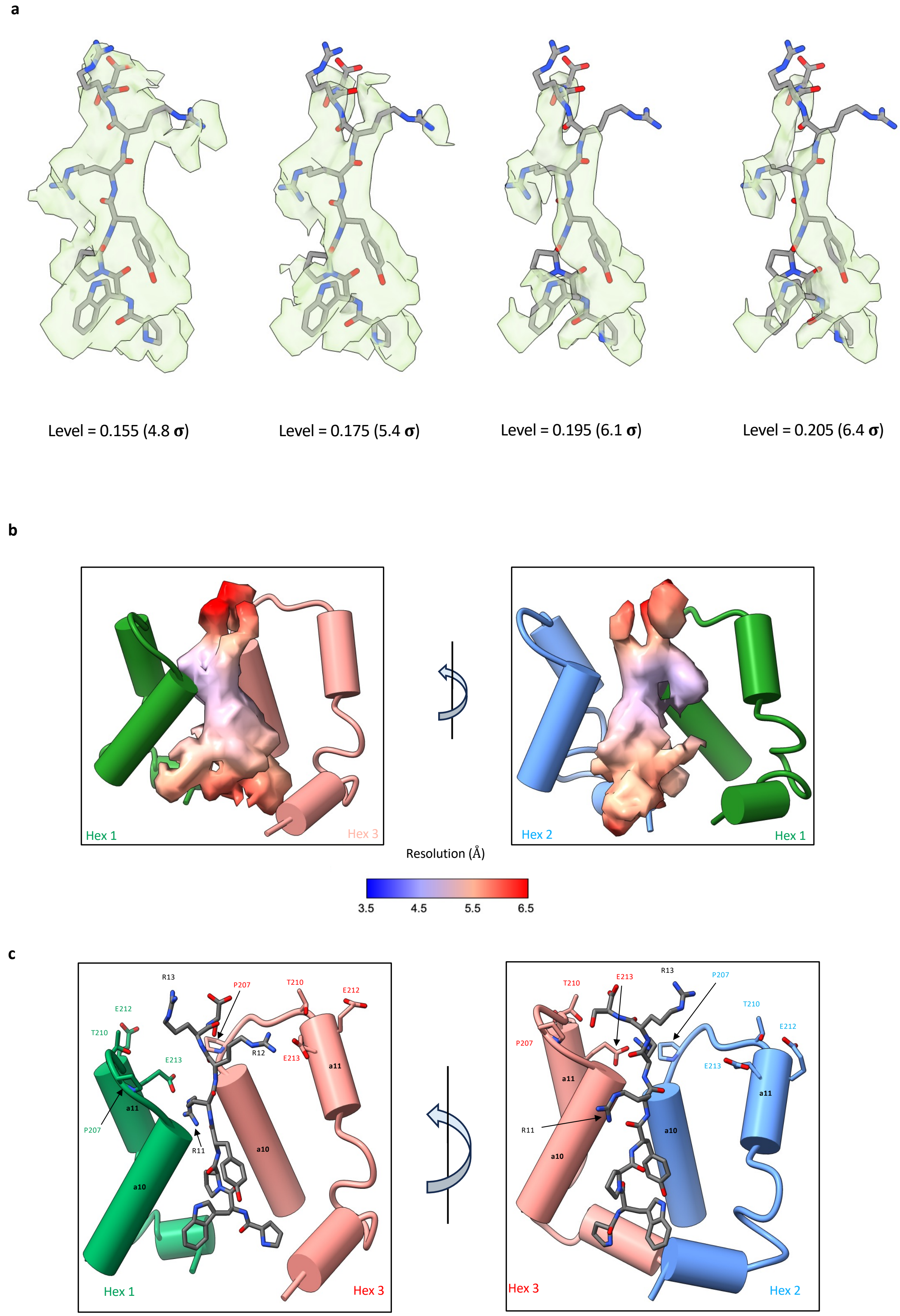


**Extended Data Figure 9: Modelling of MX2 residues 7-14**

**a,** Model of MX2 residues 7-14 within the extra density at the tri-hexamer interface of the MX2$_{1-45}$-GST S28D bound capsid map. The map has been displayed at various contour levels. **b,** Model of CA residues 170-231 (green, blue and salmon) at the tri-hexamer interface showing the density corresponding to MX2 residues 7-14 coloured by local resolution estimation. **c,** Same model as in **b**, with capsid residues harbouring known MX2 escape mutations labelled.

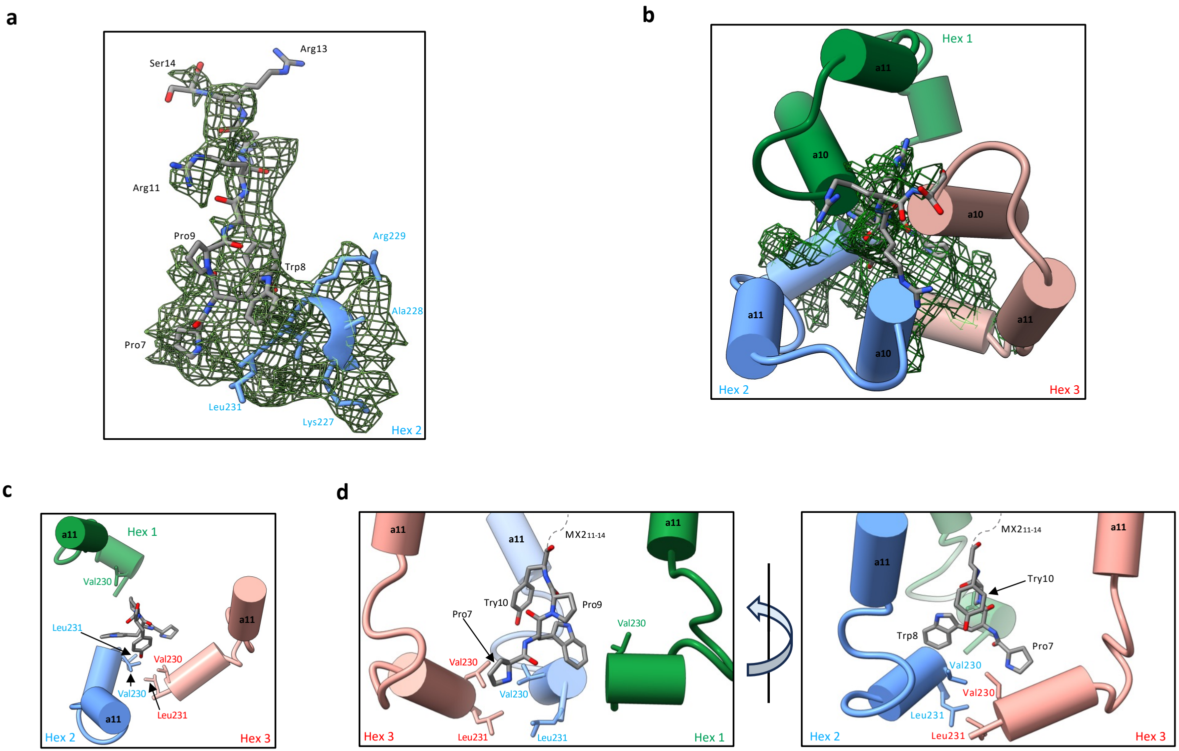


**Extended data Figure 10: MX2 promotes CA C-terminal intra-hexamer interactions**

**a,** Cryo-EM difference map. The green mesh represents the positive difference density obtained by subtracting the $MX2_{1-45}$-GST S28D bound capsid map from the unbound capsid map. The atomic models of MX2 residues 7-14 and CA hexamer 2 (blue) are shown. **b,** Top view of **a,** with CA residues 198-231 from all three CA hexamers (green, blue and salmon) composing the tri-hexamer interface shown. **c-d,** Model of the MX2 bound capsid C-terminal interactions. For clarity, only MX2 residues 7-10 (grey) and CA residues 212-231 (green, blue and salmon) are shown.
